# Proximity to Boundaries Reveals Spatial Context Representation in Human Hippocampal CA1

**DOI:** 10.1101/2023.04.18.536607

**Authors:** Maya Geva-Sagiv, Halle R. Dimsdale-Zucker, Ashley B. Williams, Charan Ranganath

## Abstract

Recollection of real-world events is often accompanied by a sense of being in the place where the event transpired. Convergent evidence suggests the hippocampus plays a key role in supporting episodic memory by associating information with the time and place it was originally encountered. This representation is reinstated during memory retrieval. However, little is known about the roles of different subfields of the human hippocampus in this process. Research in humans and non-human animal models have suggested that spatial environmental boundaries have a powerful influence on spatial and episodic memory, as well as hippocampal representations of contexts and events. Here, we used high-resolution fMRI to investigate how boundaries influence hippocampal activity patterns during the recollection of objects encountered in different spatial contexts. During the encoding phase, participants viewed objects once in a naturalistic virtual reality task, in which they passively explored two rooms in one of two houses. Following the encoding phase, participants were scanned while they recollected items in the absence of any spatial contextual information. Our behavioral results demonstrated that spatial context memory was enhanced for objects encountered near a boundary. Activity patterns in CA1 carried information about the spatial context associated with each of these boundary items. Exploratory analyses revealed that memory for the room in which each object was studied was correlated with the fidelity of retrieved spatial context representations in anterior parahippocampal cortex and subiculum. Our results highlight the privileged role of boundaries in CA1 and suggest more generally a close relationship between memory for spatial contexts and representations in the hippocampus and parahippocampal region.

## Introduction

Episodic memory, the ability to remember past events, is intertwined with the place and time (spatiotemporal context) in which an event unfolded. Indeed, Tulving (1972) argued that the memory of an event is organized with respect to its spatiotemporal relation to other events. O’Keefe and Nadel (1978) built on this idea, stating that a specific brain area, the hippocampus, supports memory for items or events within a spatiotemporal context. Consistent with these general ideas, behavioral research has shown that spatial and temporal context reinstatement can influence episodic memory retrieval (Eichenbaum, 2017; Smith and Vela, 2001), and neuroscience research has suggested that putative hippocampal representations for spatial (Chadwick et al., 2011; Deuker et al., 2016; Kyle et al., 2015; Miller et al., 2013; Nielson et al., 2015; Rosenbaum et al., 2004; Viskontas et al., 2009; Zeidman and Maguire, 2016) or temporal context (Dimsdale-Zucker et al., 2018; DuBrow and Davachi, 2016; Eichenbaum et al., 2007; Molitor et al., 2021) are reinstated during episodic memory retrieval.

Mental representations of spatial context are structured by boundaries (Epstein et al., 2017). Research in animal models and humans alike has pointed to a privileged representation of spatial boundaries in neural activity, as they play a crucial part in models of navigation (Epstein et al., 2017; Julian et al., 2018b). For example, electrophysiological recordings in rodents have identified single cells in the entorhinal cortex and the subiculum that signal location and distance from borders in explored environments (Barry et al., 2006; Lever et al., 2009; Savelli et al., 2008; Solstad et al., 2008). Consistent with these findings, hippocampal activity in humans during scene imagination relates to the number of boundaries in the environment (Bird et al., 2010), and hippocampal activity during navigation in a virtual-reality task predicts learning of object locations relative to boundaries (Doeller et al., 2008).

Converging evidence points to the importance of non-spatial event boundaries, moments in time identified by observers as a transition between events, for parsing continuous experiences (Kurby and Zacks, 2008). These transitions have been suggested to have a key role in organizing information in memory, and multiple studies have demonstrated the sensitivity and specificity of hippocampal responses to event boundaries (Baldassano et al., 2017; Ben-Yakov and Henson, 2018; Cohn-Sheehy et al., 2021; Reagh et al., 2020; Yoo et al., 2022; Zheng et al., 2022).

Here, we used a virtual reality exploration task paired with high-resolution fMRI to investigate how boundaries are reflected in the neural activity of the hippocampus and the extended episodic memory network (Maguire et al., 1998), consistent with the idea that the hippocampus binds information about space and time to create a unified representation of an experienced event (Eichenbaum et al., 2007; O’Keefe and Nadel, 1978) and that this representation is reinstated during memory retrieval.

The hippocampus is composed of multiple interconnected subfields that contribute to memory processing and representation (Amaral and Lavenex, 2007). Characterizing their roles and interplay is essential for understanding how the hippocampus encodes context representations, which are the building blocks of complex episodic memories (Eichenbaum et al., 2007; O’Keefe and Nadel, 1978). Computational models propose that the unique anatomical properties of dentate gyrus and CA3 make these subfields ideal for pattern separation, the orthogonalization of highly similar inputs though sparse firing, whereas the CA1 and subiculum subfields are more suited to pattern completion, i.e. preserving features that are common to different inputs (Guzowski et al., 2004; Marr, 1971; Norman and O’Reilly, 2003; Schapiro et al., 2017; Treves and Rolls, 1994; Yassa and Stark, 2011; Y. Zheng et al., 2021). It is unclear, however, whether these concepts capture the functional differences between subfields of the human hippocampus.

One method that has been used to uncover differences between hippocampal subfield representations is to compare their activity based on high-resolution functional magnetic resonance brain imaging (fMRI) measurement while subjects are asked to identify previously memorized visual stimuli and reject highly similar lures (Bakker et al., 2008; Berron et al., 2016; Lacy et al., 2010; Stevenson et al., 2020). Brain activity in these studies has been interpreted to reflect a bias towards pattern completion in CA1 and a tendency for pattern separation in CA3/DG. Can these results, based on unique visual stimuli, be extended to the prediction of hippocampal subfields code of spatial context? In a study by Kyle (Kyle et al., 2015), virtual environments were used to test context-specific encoding in hippocampal subfields. Participants were first acquainted with four spatial contexts in a virtual environment and were later asked to estimate spatial distance between locations based on their acquired spatial knowledge. Comparing multivariate response patterns during these judgments demonstrated unique activation (remapping of representations) between spatial contexts in CA3/DG and less so in CA1. Pattern separation in CA3/DG of different spatial contexts was also demonstrated by Zheng (L. Zheng et al., 2021), while others reported increased overlap for CA3 for similar environments (Stokes et al., 2015) and reduced overlap for CA1 (L. Zheng et al., 2021). These conflicting results point to the need for a deeper understanding of the interplay between subfields in the hippocampal circuit with respect to space. To our knowledge, no previous high-resolution fMRI studies have reported incidental reinstatement of spatial context information in particular hippocampal subfields during episodic recollection (without explicit retrieval of spatial information during scans).

In the present study, we revisited the data from Dimsdale-Zucker et al, 2018 (<u>OSF | abcdcon_pub</u>) to gain further insight into the nature of spatial context representation in the hippocampus in a virtual-reality paradigm that mimics the natural exploration of spatial environments. Participants viewed a set of videos, containing objects residing in two different spatial contexts (two houses), and they were scanned while they recollected these objects in isolation, devoid of any contextual information (Fig. 1A-B). Using high-resolution fMRI, we previously found that hippocampal subfields CA1 and a combined CA2/CA3/DG (CA23DG) play complementary roles in supporting the retrieval of episodic context (each video representing a unique episode) (Dimsdale-Zucker et al., 2018). In our previous study, we compared hippocampal activity pattern similarity (PS) for pairs of objects encountered in the same house (i.e., same spatial context) against PS for pairs of objects that were encountered in different houses (i.e., different spatial context). Initial analyses revealed no significant effects of spatial context in any hippocampal subfields or even in extrahippocampal regions such as the parahippocampal cortex (PHC) that are thought to support spatial context representation (Bastin et al., 2013; Brown et al., 2010; Deuker et al., 2016; Eichenbaum et al., 2007; Kunz et al., 2021; Morgan et al., 2011). This null finding was somewhat surprising, given that participants were able to recall spatial contextual information about many of the studied objects.

**Figure 1:**
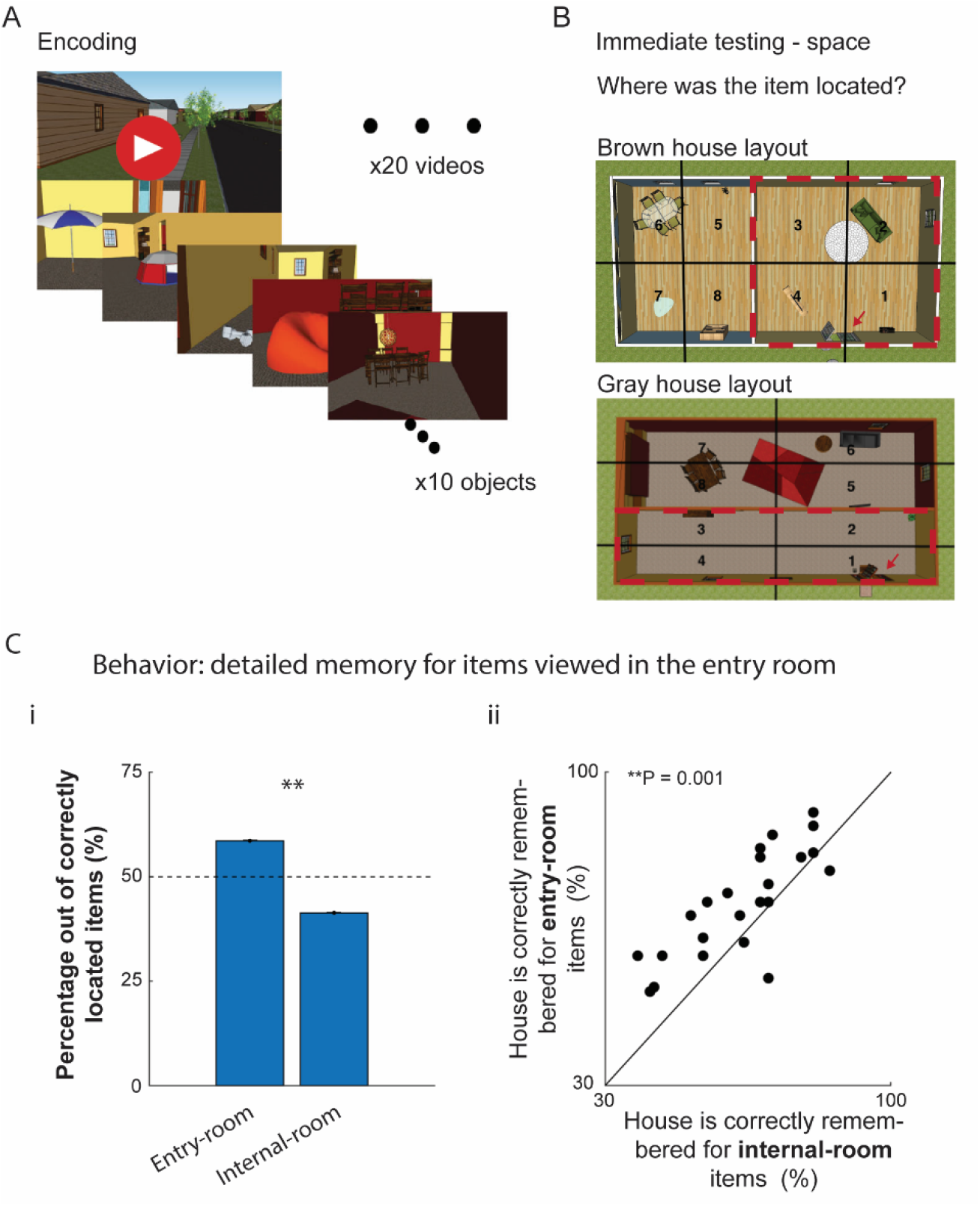
Revealing spatial context in a virtual-reality paradigm. (A) Experimental design: Participants encoded objects uniquely located within one of two houses (spatial contexts) across a series of 20 videos (episodic context). Each video was seen once (one-shot learning). (B) At the end of each video, participants were asked to place the encoded objects on a bird’s eye map. Note that the two spatial contexts differed by their aspect ratio. The magenta arrows mark the entry doors’ locations and the magenta dashed line marks the entry room of each house. (C) Behavior: (i) Encoding of objects viewed in the entry room was significantly better than that of objects encountered in the internal room: Each bar depicts the average number of successfully-placed images from each room, which is expected to be 50% if room identity does not affect memory; (ii) In a spatial memory test, taken after scanning, subjects were significantly better in recalling the house an object was viewed in for entry-room objects. Each dot represents one subject, y-axis represents the percentage of correct house choices for entry-room objects and x-axis is for internal room objects. (** denote P < 0.01, using a paired, two-sided Wilcoxson ranksum test).

In our re-analyses, we considered several possibilities: First, we investigated whether spatial context representation in hippocampal subfields might be tied to the successful encoding of the spatial context. We, therefore, examined pattern-similarity (PS) during retrieval specifically for objects that subjects correctly associated with the specific room in which objects were encountered during encoding. Second, we considered whether the hippocampus might particularly carry information about objects viewed in the entry room of each house. These objects might be especially significant, given evidence for enhanced hippocampal encoding during event boundaries (Ben-Yakov and Henson, 2018; Cohn-Sheehy et al., 2021; DuBrow and Davachi, 2016; Reagh et al., 2020; Swallow et al., 2011) and hippocampal sensitivity to spatial boundaries (Bird et al., 2010; Horner et al., 2016; Julian et al., 2018a, 2018b; Lever et al., 2009; Stangl et al., 2021). In our experimental paradigm, objects encountered in the entry room were both in proximity to the event boundary (start and end of the virtual tour) and near the boundary of the spatial context (entry point of the house). These considerations, as well as behavioral data in our study, suggesting that subjects generate a more detailed memory of these objects, led us to hypothesize the hippocampus would reflect a dedicated code for these entry room items.

We also considered the possibility that areas in the navigation network (Maguire et al., 1998) outside of the traditional hippocampal circuit might carry information about the spatial context. We focus here on the subiculum and the parahippocampal cortex (PHC) (Aggleton et al., 2010; Epstein, 2008; Kravitz et al., 2011; Ranganath and Ritchey, 2012) that encode information about spatial (Bastin et al., 2013; Brown et al., 2010; Deuker et al., 2016; Eichenbaum et al., 2007; Kunz et al., 2021; Morgan et al., 2011) and temporal context that is reinstated during recollection (Diana et al., 2013; Düzel et al., 2003; Essoe et al., 2022; Khader et al., 2005; Rugg and Vilberg, 2013). Recollection-related activity in PHC has been interpreted as having a central role in the representation of contextual information (Diana et al., 2013; Rugg and Vilberg, 2013). Subiculum, an output structure of the hippocampal circuit (Amaral and Lavenex, 2007; Ding, 2013), has also been linked to memory for spatial context (Lever et al., 2009; Peer et al., 2019; Zeidman et al., 2015; Zeidman and Maguire, 2016). Both the PHC and subiculum have been demonstrated to be reliably recruited during virtual spatial navigation in humans (Brown et al., 2010; Spreng et al., 2009; Zeidman et al., 2015). We hypothesized that better recollection performance would predict a more detailed memory for objects seen during the encoding phase, and tested for relationships between spatial context representations and individual differences in memory accuracy in these regions. We report here how spatial context, which is incidental to the recollection task, is reflected in hippocampus, subiculum, and PHC brain activity during successful recollection.

## Results

### Behavioral Results – Enhanced source memory for objects in the entry room

To explore spatial context reinstatement in hippocampal subfields, participants were introduced to a virtual reality environment consisting of two houses, each house containing two rooms (spatial contexts; Fig. 1A-B). After becoming familiarized with the appearance and spatial layouts of each house, which differed in their color and rooms’ aspect ratio (but were equal in overall exploration space, Fig. 1B), participants viewed a series of 10 videos presenting first-person navigation through each house, encountering a series of 10 unique objects in each video (Fig. 1A). The virtual tour passively led viewers through the houses, starting with an entry room, continuing to an internal room, and backtracking to exit the house through the entry room.

Subjects’ memory performance was evaluated during three phases of the experiment – during encoding, in the fMRI scanner, and during a final post-scan test. During encoding, at the end of each video viewing, participants were asked to judge objects’ value and to place the objects they saw in the video on a bird’s eye map of the house (Fig 1B). Note that each video was viewed once and no feedback was given on the placement test. Following this study phase, participants performed an object recognition test that required them to differentiate between studied and novel objects, while undergoing fMRI scanning. Critically, during the scanned object recognition phase, objects were presented in isolation devoid of any explicit contextual information from encoding. Lastly, back in the lab, in a third testing phase which was performed immediately after MRI scanning, subjects were asked to recall where (house and room) every object had been studied.

Behavioral results have been reported extensively in Dimsdale-Zucker et al. (Dimsdale-Zucker et al., 2018), so we will only briefly summarize here the key findings relevant to the current re-analysis. First, at the end of each video viewing, during the encoding phase, subjects indicated the correct room for each object with very high probability (correct room placement rate = 0.96±0.02 (mean ± std), 75% of the objects viewed were correctly placed in the correct room by all participants in this immediate test, Sup Fig. 2A). Second, recognition memory performance was measured by evaluating responses to new and old objects during fMRI scanning. Hit-rate was high with some variability between subjects’ performance (“remember” hit rate = 0.66± 0.17 (mean ± std), Sup Fig. 2B, (Dimsdale-Zucker et al., 2018)).

Based on previous studies reporting enhanced recognition of objects that were presented around event boundaries (Newtson and Engquist, 1976; Schwan et al., 2000), we compared memory performance for objects presented in the entry room against objects presented in the internal room of the house. Analyses of the behavioral data revealed superior performance for objects residing in the entry room in three separate phases of testing: (i) At the end of each video viewing, during the encoding session, participants placed objects they had seen in the entry room with higher accuracy than objects viewed in the internal room (Fig 1C, paired Wilcoxon test, P < 10^-4^). (ii) During MRI scanning, when participants were asked to recognize previously learned objects, objects in the entry room had a higher (though non-significant) probability of being recollected relative to objects located in the internal room (paired Wilcoxon test, P = 0.06). (iii) In the last testing phase, following the MRI scan, subjects’ memory of the house the object was viewed in was significantly higher for entry room objects (Wilcoxson signed-rank test, P = 0.0014, Fig 1Cii).

We considered the possibility that recency or primacy effects (Ebbinghaus, 1885) may explain the superior accuracy when recalling the location of objects in the entry room, relative to internal room ones, and whether the superiority was strictly found in the temporal aspect of the memory (order of events), rather than a better spatial source memory. As room identity overlaps with the order of viewing of objects within a video clip (i.e. a subject can infer the room location based on the time an object was seen), these effects could indeed underlie the superior memory of entry room objects in the encoding and MRI test phases. However, the house identity cannot be inferred from the viewing order, so superior memory for entry-room objects’ house identity (Fig. 1Cii), suggests a better spatial context memory was acquired for entry-room objects relative to the inner room ones. Taken together, these behavioral results suggest that subjects acquired a more comprehensive memory of entry-room objects, relative to the inner-room objects, and that the privileged memory status of entry-room objects persisted from the encoding phase to the last phase of testing (a couple of hours later).

### Proximity to boundaries shapes neural overlap in hippocampal subfields

We expected recollection-based item recognition to trigger reactivation of information about the context in which that object was encountered (Deuker et al., 2013; DuBrow and Davachi, 2016; Eichenbaum et al., 2007; Miller et al., 2013; Nielson et al., 2015). Accordingly, we tested whether multi-voxel patterns elicited during object recollection, presented without any house/video information, carried information about the associated spatial location for each object. To do this, we used Representational Similarity Analysis (RSA) (Dimsdale-Zucker and Ranganath, 2018; Kriegeskorte et al., 2008b) to measure the similarity of fMRI multi-voxel patterns.

We estimated single-trial multi-voxel patterns within regions of interest corresponding to CA1 and a combined CA2/CA3/dentate gyrus (CA23DG) subregion within the body of hippocampus (Methods). We then computed voxel pattern similarity (PS) between trial pairs for successfully recollected objects. Having discovered that spatial memory accuracy was behaviorally enhanced for entry room objects relative to the internal room ones (Fig. 1C), we tested whether hippocampal activity patterns might disproportionately carry information about the location of objects encountered in the entry room, as compared with objects from the inner room. We restricted the analysis to object recognition trials that were associated with accurate spatial memory (i.e., objects placed in the correct house and room) during the encoding phase (Sup. Fig. 2A). We calculated PS for pairs of objects that were seen in the same house, separately for pairs of objects encountered in the entry room, pairs of objects encountered in the inner room, or pairs that were encountered in different rooms of the same house. To control for the possibility that any observed effects are attributed to temporal context (these effects were previously reported in (Dimsdale-Zucker et al., 2018), we eliminated trial pairs that had been studied within the same video to ensure that any observed effects could uniquely be attributed to spatial context.

To test whether hippocampal subfields carried information about an object’s spatial location, we fitted a linear mixed effect model with a random effect of Subject (Gordon et al., 2014) testing for effects of ROI (CA1 and CA23DG), Spatial location in the house (entry room vs. inner room vs. different room) as well as their interactions on multivoxel PS (Methods). Adding hemisphere as another factor did not reveal a significant interaction so the reported analyses are aggregated across the two hemispheres. We found a highly significant interaction between ROI x Spatial location in the house (χ^2^(2) = 25.54, P < 10^-4^).

To unpack this interaction, we used a mixed effects model to test effects for item-pairs’ PS as a function of Spatial Location (same house: entry room vs. inner room vs. different room) separately for each ROI. For both CA1 and CA23DG, there was a significant effect of Spatial location in the house^1^ (Fig. 2Bi^2^, Mixed effects model, CA1 – χ^2^ (2) = 14.39, = P = 0.0007; CA23DG – χ^2^(2) = 7.54, P = 0.02). In CA1, activity patterns were more similar for item pairs seen in the entry room than for objects seen in the inner room (χ^2^ (1) = 13.70, P = 0.0002) and for objects seen in different rooms (χ^2^ (1) = 13.70, P = 0.003). These results are consistent with the idea that activity patterns in CA1 carry information about the entry room as a spatial context. In contrast, in CA23DG, pattern similarity was higher for inner room pairs than entry room pairs, (χ^2^ (1) = 7.27, P = 0.006), but there was no significant difference between inner room pairs and pairs of objects encoded in different rooms (χ^2^ (1) = 1.4, P = 0.22). Although these results indicate that CA23DG pattern similarity was sensitive to the particular room that an object was encountered in, there was no strong evidence for a spatial context representation per se, because neither entry room nor inner room pairs differed from across-room pairs.

**Figure 2:**
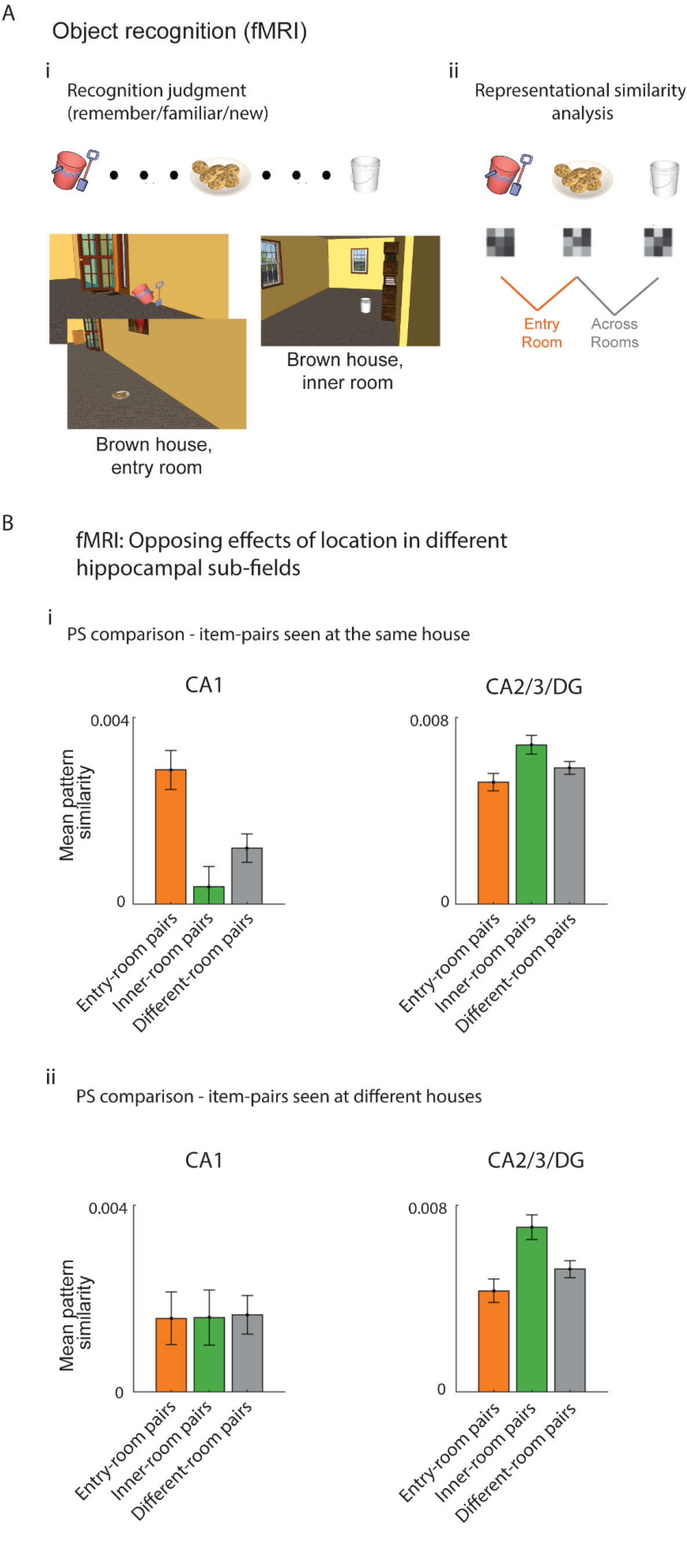
Pattern similarity in CA1 and CA23DG reflects proximity to spatial and temporal boundaries. (A) (i) Experimental design: Subjects were scanned (high-resolution fMRI) while performing an object recognition test. Objects were presented without any contextual information. The images below depict the locations the example objects were seen during the encoding session. (ii) Representational similarity analyses (RSA) was used to compare multi-voxel patterns elicited by each recollected object relative to other recollected objects based on their spatial context. (B) fMRI: (i) Pattern similarity was calculated for regions of interest corresponding to CA1 and a combined CA2/CA3/dentate gyrus (CA23DG) subregion within the body of hippocampus (Methods). Same house pairs: Pattern similarity was highest in CA1 subfield for object pairs from the same house when located in the entry room relative to inner-room pairs and to pairs of objects viewed in different rooms. CA23DG showed a different pattern such that pattern similarity was lower for object pairs seen in the entry room of the same house relative to inner room pairs. (ii) Different house pairs: CA23DG - pattern similarity was lower for object pairs when both were seen in the entry room (close to the event boundary) of different houses relative to inner room objects, while no significant difference was found in CA1. The effect in CA23DG could stem from reduced overlap for entry-room pairs or increased overlap between inner-room pairs which was higher than overlap between different-room pairs.

We next investigated the possibility that subfields might encode a general representation of information encoded close to a boundary. For instance, it is conceivable that the hippocampus could encode objects as having been in an entry room, as opposed to encoding a representation of an object that was encountered in a specific room of a specific house. To test this possibility, we performed the same analyses described above but restricted our analyses to objects encountered in different houses. Interestingly, this analysis revealed a significant effect of Spatial location (different house: entry room vs. inner room vs. different room) in the house for CA23DG (χ^2^ (2) = 12.67, P = 0.001), but not in CA1 (χ^2^ (2) = 0.06, P = 0.9; see Fig. 2Bii^2^). Follow-up tests to break down the effect of Spatial Location in CA23DG revealed that pattern similarity was higher for inner room pairs than entry room (χ^2^ (1) = 12.68, P = 0.0003) or different room (χ^2^ (1) =6.96, P = 0.08) pairs.

Our data suggest that CA1 and CA23DG represented different aspects of the room contexts. CA1 activity patterns were consistent with a privileged representation of the entry room within each house. CA23DG activity patterns, in contrast, seemed to reflect information common to any object in an inner room.

### Exploratory Analysis - neural spatial sensitivity in aPHC and subiculum

Dimsdale-Zucker et al. (Dimsdale-Zucker et al., 2018) focused on results from CA1 and CA23DG, leaving open the question of whether spatial context information might be incidentally reinstated in other MTL subregions that have been associated with successful recollection. To address this question, we ran a series of exploratory, post-hoc analyses to develop hypotheses that can be tested in future studies. We focused our analysis on two subregions that have been implicated in episodic memory and navigation - the anterior parahippocampal cortex (aPHC) (Baldassano et al., 2016; Bar et al., 2008; Diana et al., 2007; Eichenbaum et al., 2007; Epstein, 2008; Epstein and Baker, 2019; Ozdogan and Morris, 2012; Suzuki and Amaral, 2004) and the subiculum (Amaral and Lavenex, 2007) (see Supplementary Fig. 1, see Methods for segmentation process).

We first examined whether, as in the hippocampus, aPHC and subiculum might carry information about entry- or inner-room objects. A mixed effects model revealed no effects of spatial location, suggesting that there was no significant sensitivity to proximity to spatial or event boundaries as we found in hippocampal subfields. We then examined whether voxel pattern information in these areas might carry spatial information at a more coarse level, reflecting the recollected spatial context (house) associated with each object. To address this question, we computed a ‘spatial sensitivity score’ by calculating the difference between the median PS values of objects located in the same room of the same house, relative to objects from different rooms in different houses (Fig. 1Cii). However, when aggregating trials from all subjects, we did not find a significant spatial sensitivity score in aPHC nor subiculum.

We next explored the possibility that this null finding was due to interindividual differences in spatial memory accuracy (Supplementary Fig. 2). Consistent with this hypothesis, we found a significant correlation between the ‘spatial sensitivity score’ and recollection success (hit-rate) per subject in both right hemisphere’s aPHC (Pearson correlation: R=0.53, p=0.008, Fig 3i) and subiculum (Methods, Pearson correlation: R=0.54/p=0.007, Fig3ii). We did not observe a significant correlation for responses in the left hemisphere for either region. Overall, results from these exploratory analyses suggest that aPHC and subiculum carry information about recollected spatial contexts at a relatively coarse level and that these representations were tied to higher spatial memory accuracy.

**Figure 3:**
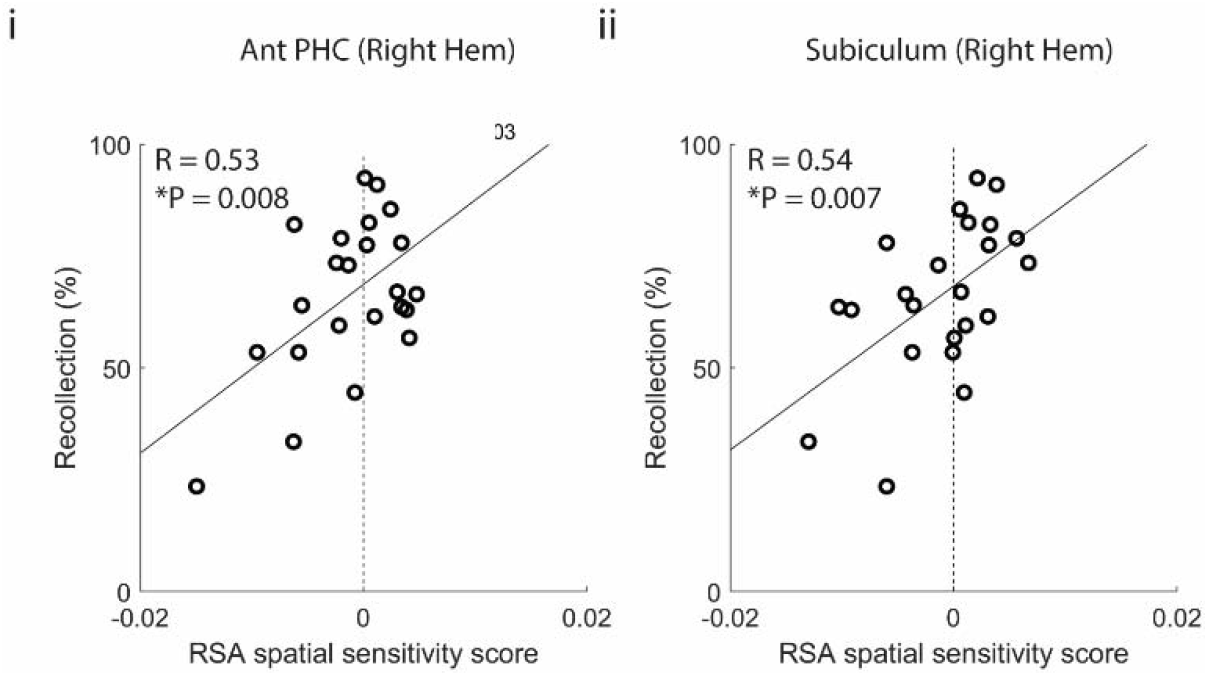
‘Spatial sensitivity score’ of aPHC and Subiculum is tightly correlated to recollection performance during fMRI scan. Spatial sensitivity score was defined as the difference between medians of pattern similarity values for objects from the same house and room vs different rooms in different houses (the largest spatial difference obtained in the experienced virtual-reality space). We found a tight correlation between this measure and recollection success (hit-rate) per subject – recollection success was a predictor of spatial sensitivity as reflected by neural pattern similarity in anterior PHC (i) and subiculum (ii) in the right hemisphere. Each circle represents one subject, R and P values for Pearson correlation are calculated over all subjects (n = 23).

## Discussion

Boundaries are thought to be useful to parse continuous experiences into meaningful chunks in space and time (Kurby and Zacks, 2008; Lu et al., 2022), and accumulating evidence suggests hippocampal activity is enhanced during the perception of boundaries in naturalistic events (Baldassano et al., 2017; Ben-Yakov and Henson, 2018; Cohn-Sheehy et al., 2021; Reagh et al., 2020; Yoo et al., 2022). Spatial boundaries are crucial for navigation and are over-represented by neural activity during active and virtual behavior (Barry et al., 2006; Bird et al., 2010; Doeller et al., 2008; Julian et al., 2018a, 2018b; Lever et al., 2009; Savelli et al., 2008; Solstad et al., 2008). Here, simulating the exploration of two spaces, combined with object recognition testing collected during high-resolution fMRI scanning, we found that spatial context memory was enhanced for objects in proximity to a boundary and that CA23DG and CA1 showed qualitatively different representations of retrieved spatial context for these objects. Our results highlight the privileged role of boundaries in the representation of continuous events in memory and suggest that while CA1 is more informative regarding spatial context per-se, CA23DG is more informative with respect to event boundaries.

Dimsdale-Zucker et al. (2018) previously demonstrated that CA1 exhibited increased representational overlap for objects sharing an episodic context (i.e., objects seen in the same video), while CA23DG exhibited greater PS for objects from distinct as compared to shared episodic contexts. However, analyses reported in Dimsdale-Zucker et al. (2018) revealed no evidence of spatial context representation in these subfields. The present analyses were designed to more precisely characterize spatial context representation in the hippocampus in the following way: First, we restricted our analysis to objects for which subjects demonstrated accurate spatial memory. Second, we only calculated PS across pairs of objects seen in different videos to identify spatial context representations in the absence of any shared episodic context. Third, and most importantly, we differentiated between activity elicited by objects that were encountered in the entry room, which was both an environmental boundary and an episodic event boundary in these videos, and the inner room of each house. The latter factor was critical, as we identified a spatial memory advantage for objects encountered in the entry room, as well as the reinstatement of the entry-room context in CA1 during the recollection of studied objects.

The present results are broadly consistent with evidence from both animal models and human studies suggesting that environmental boundaries play a crucial part in models of navigation (Julian et al., 2018b). In rodents, studies have found cells that signal borders in subiculum and medial entorhinal cortex (Hartley et al., 2014; Lever et al., 2009; Solstad et al., 2008). In humans, environmental boundaries serve as a primary cue for reorientation when one loses their bearing (Cheng et al., 2013; Epstein et al., 2017) and boundary-related neural signatures have been observed using fMRI and intracranial electroencephalographic (iEEG) recordings (Bird et al., 2010; Doeller et al., 2008; Horner et al., 2016; Julian et al., 2018a; Lee et al., 2018; Shine et al., 2019; Stangl et al., 2021). An interaction between spatial boundaries in virtual-reality tasks and long-term memory has been shown before to negatively impact memory retrieval when boundaries break the encoding process (Horner et al., 2016; Radvansky et al., 2011).

The present study differs from many of the studies described above in two ways. First, we did not directly assess spatial memory or navigation, but, instead, examined incidental retrieval of spatial context representations during an object-recognition test. Thus, our study was intended to assess how recollection of previously encountered objects leads to recovery of associated contextual information (Diana et al., 2007; Eichenbaum et al., 2007). The second key difference was that each spatial context (i.e., each house) consisted of two rooms, whereas previous studies, to our knowledge, examined activity associated with open environments (Deuker et al., 2016; Kyle et al., 2015; Peer and Epstein, 2021; Steemers et al., 2016; Stokes et al., 2015; L. Zheng et al., 2021), single rooms (Guo et al., 2021), or adjoining rooms within a single context (Hassabis et al., 2009; Kim and Maguire, 2018). This distinction turned out to be important because PS values in CA1 were greater for pairs of objects that were located in the entry room of the same house than for pairs of objects in the inner room or objects encountered in different rooms. This finding only applied to objects that were in the same house— objects that were in the entry rooms of different houses did not show pattern similarity increases. Accordingly, these results suggest that, during recollection, CA1 patterns carried information about the specific spatial context in which entry room objects were encountered.

One potential limitation of the present study concerns the relationship between the entry room context and the temporal position of objects in each video. The results observed in CA1 cannot be attributed to reinstatement of a global episodic context, because all of the present analyses excluded trial pairs corresponding to objects from the same movie. However, because entry room objects were encountered at the beginning and end of each movie, proximity to the entry room was related to proximity to a temporal/event boundary as well as the spatial boundary for the entry and exit of the house. Previous studies have demonstrated that objects near the start and end of videos (which can be considered temporal event boundaries) tend to be better remembered (Newtson and Engquist, 1976; Schwan et al., 2000). For instance, Horner et al (2016) reported that a sequence of two objects presented in the same room in a virtual reality environment is more accurately remembered than a sequence of two objects presented in adjoining rooms, suggesting that the presence of a spatial boundary at encoding (a doorway between two rooms) impairs subsequent recollection of the order that objects were presented in. In the present study, objects were encountered in the context of short videos that simulated a walk through a house with one main door.

It is conceivable that participants could have used temporal boundaries as an anchor point, such that an object is simply remembered as simply being at the beginning or end of a movie. That said, our results were not consistent with the idea that CA1 representations were related to proximity to an abstract event boundary (Kurby and Zacks, 2008). Pattern similarity in CA1 was increased for entry room objects from the same house, but no significant effect was observed when comparing entry room objects from separate houses. Thus, our results suggest that the CA1 representations of entry room objects had a spatial component.

Results in CA23DG were qualitatively different from those in CA1. Unlike CA1, we did not see any evidence for a significant spatial context representation of the entry room, and PS for entry room objects was lower than for inner room objects (Fig. 2B, right panels). The reduction in pattern similarity between entry-room objects relative to inner-room pairs was evident regardless of whether the objects were in the same or different houses. There are at least two ways to explain this pattern of results. It is possible that CA23DG representations of objects that appeared in an entry room (of any house) were pulled apart from one another over the course of learning, consistent with prior reports of hippocampal “repulsion” (Chanales et al., 2017; Ritvo et al., 2019) in which neural representations of overlapping memories are hyper-differentiated from one another. Alternatively, it is possible that CA23DG preferentially encoded a common contextual representation for any object that was encountered in an inner room (i.e., far from an event boundary). The present results do not allow us to make strong interpretations in favor of either account. That said, the different patterns of results in CA1, as compared with CA23DG, suggest the need for future research investigating the extent to which spatial representations may be shared, hyper-differentiated, or abstracted.

In addition to investigating activity in CA1 and CA23DG, we also conducted exploratory analyses of spatial representations in aPHC and subiculum. These areas are major input and output hubs to the hippocampus (Aggleton et al., 2010; Amaral and Lavenex, 2007; Burwell, 2000; Epstein, 2008; Kravitz et al., 2011; Ranganath and Ritchey, 2012; van Strien et al., 2009), and known to play an important role in spatial and episodic memory (Eichenbaum et al., 2007; Ekstrom and Ranganath, 2018; O’Mara et al., 2009). For example, Hassabis et al (2009) examined activation while participants navigated a virtual environment consisting of two connected square rooms and found that activation patterns in parahippocampal cortex distinguished between the rooms. To identify whether fMRI responses in our study reflect the similarity of locations in the virtual space (defined by house and room identity), we compared PS overlap for item pairs from same house/room vs item pairs from different houses/rooms. Interestingly, we observed a robust correlation in these areas between individual hit-rate in the recognition task, and a difference in PS values for item pairs from same/different spatial contexts. It is important to reiterate that these analyses in subiculum and aPHC were post-hoc and exploratory, but the results are sufficient to motivate future work to characterize spatial and episodic memory representations in the broader parahippocampal region (Witter et al., 1989).

One of the hallmarks of episodic memory is the ability to bring to mind a coherent representation of the elements of an event, even when these elements are incidental to the retrieval condition (Tulving et al., 1983). Our findings complement previous studies that revealed hippocampal reinstatement of neural activity during a recognition task in intracranial patients (Miller et al., 2013) and in fMRI studies (Chadwick et al., 2011; Kyle et al., 2015; Zeidman and Maguire, 2016; L. Zheng et al., 2021) during tasks that involve intentional recollection of the spatial locations in which objects were studied. Our results indicate that, during recollection, just as Tulving posited, spatial information is retrieved and reflected both in MTL subregions (aPHC and subiculum) and hippocampal subfields (CA1 and CA23DG), even when incidental to the recognition task.

## Methods

### Participants

Analyses presented are from twenty-three participants (N_female_ = 11, N_male_ = 22 based on self-reporting, mean age = 19.5 years), the same cohort described in Dimsdale-Zucker (2018).

### Stimuli and materials

Study materials included two virtual houses, each divided into two rooms, which were created in Google SketchUp (https://www.sketchup.com/, version 15.3.329, all stimuli are freely available – <u>OSF | abcdcon_pub</u>, https://osf.io/5th8r/). The two houses differed in their exterior color, wall color, room orientation, and decoration style (Fig. 1B). Though they differed in their aspect-ratio houses were matched in their total virtual square area. Each house contained ten pieces of landmark furniture that shared semantic labels (e.g., “couch”) but differed in appearance (e.g., angular gray couch vs. plush green couch). Three hundred neutral objects (e.g., football helmet, suitcase, teddy bear) were selected from the Google SketchUp image library. Two-hundred and forty of these objects were randomly selected and the rest were used as lure objects (see below). Object assignment was random and was house- and video-unique so that 12 lists, each consisting of 10 objects, were assigned to each house. To determine object placement, rooms were divided into eighths, all possible combinations of five positions were generated, and objects were randomly assigned to one of these combinations. Thus, object configurations within each room (and thus within each house) were video-unique. Videos depicting trajectories through the houses were generated using Google SketchUp’s animation feature. Videos were exported for each house both with landmark furniture only and for each of the 12 videos with objects within each home. Trajectories did not change between videos within a home and followed a path that started at the main door circling the entry room, then crossing another door to the internal room, and backtracking to exit via the main door. Each video was approximately 1 min 40 s in duration.

### Experimental procedure

After providing informed consent, participants completed four experimental phases which were reported previously (Dimsdale-Zucker et al., 2018) and summarized here:

i. Spatial context familiarization: Participants were acquainted with two different houses and were then given up to 10 min to draw a map of each house to ensure thorough knowledge of spatial layouts (including the location of doors, walls, and furniture). Their drawings were reviewed by the experimenter to ensure any mistakes (although rare) were corrected.
ii. Object encoding: During this phase, participants viewed a series of 20 videos, each depicting passive navigation through one of the two spatial contexts (brown or gray house, Fig. 1B), with ten different objects placed along the pre-defined trajectory. At the end of each video, a still frame of each of the ten objects viewed in the house appeared one at a time in random order for 4 s while the participant judged whether the object was worth more than $50 (yes/no). After making this value judgment, the still frame of the object was replaced by a birds-eye-view perspective map of the house where each room was divided into quadrants with numeric labels. Participants indicated the quadrant number where the object had been located during the video via keypress (1–8). In total, participants saw ten videos in each house presented such that the order of the houses alternated (e.g., house1/video1, house2/video2, house1/video3, house2/video4, etc.). The ten videos were randomly selected from the pool of 12 videos in each house and the order of video presentation was uniquely randomized for each participant. The encoding phase took roughly 1 hour to complete.
iii. Object recognition test (fMRI): this phase took place across four runs in the MRI scanner following the acquisition of structural scans. Sixty-three still images were presented in each run for 3 s with a jittered inter-trial interval ranging from 2 to 8 s. All 200 studied objects were presented with an additional 52 new, unstudied objects. New objects were randomly selected from a pool of objects that were not presented in either house. While the image was on the screen, participants made recognition judgments (“remember”, “feels familiar”, “new”) via an MRI-compatible button box. Participants were instructed to make a “remember” response when they could recall a specific detail from when they had studied the object (Yonelinas, 2002), “feels familiar” if they thought they had studied the object but were unable to retrieve a specific detail and “new” for objects that they did not think they had studied during object encoding. Critically, no house or video information was re-presented to participants in these object images.
iv. Spatial context memory test: Participants were brought back to the lab to complete testing. In this phase, participants were re-presented with the 200 studied objects, and they were asked to recall where (house and room) each object had been studied. Images remained on the screen for 3 seconds while participants made their responses. There was no opportunity to skip a spatial judgment.

### fMRI acquisition and preprocessing

Extensively described in Dimsdale-Zucker (2018).

### ROI segmentation

Hippocampal subfield ROIs segmentation was described in (Dimsdale-Zucker et al., 2018) and summarized here. The ASHS toolbox (Yushkevich et al., 2010) with the UPenn atlas was used to create segmented ROIs, which were co-registered to the mean functional image and split into masks for each ROI of interest. We took a conservative segmentation approach in which we combined subfields DG, CA3, and CA2 into a combined region based on prior high-resolution fMRI work (Dimsdale-Zucker et al., 2018; Ekstrom et al., 2009; Stokes et al., 2015; Yushkevich et al., 2010). Head, body, and tail were manually defined but subfield comparisons were limited to the body where boundaries between CA1 and CA23DG can be most clearly and reliably delineated (Dimsdale-Zucker et al., 2018; Yushkevich et al., 2015a). The disappearance of the gyrus intralimbicus was used as the head/body boundary (Frankó et al., 2014) and the presence of the wing of the ambient cistern demarcated body/tail (Yushkevich et al., 2015b).

Boundaries between PRC, ERC, PHC, and subiculum regions were defined manually on individual participant high-resolution structural images in FSLView according to (Duvernoy, 1998; Insausti et al., 1998; Zeineh et al., 2001) (Supplementary Fig. 1). The PHG mask contained gray matter voxels along the banks of the collateral sulcus. Anterior PHC regions of interest (ROI) extended from the first slice immediately following the PRC, 4mm following the HC head slice to the slice where colliculi are seen (approximately 5 mm to the end of the HC, (Frankó et al., 2014)) in the anterior-posterior direction, and from the collateral sulcal fundus to the most medial vertex of the parahippocampal gyrus in the lateral– medial direction. PHC ROIs for the right and left hemispheres were created separately by manual tracing of gray matter within these demarcations.

### Data analysis

ROI summary statistics, including pattern similarity analyses, were computed using custom code implemented in MATLAB r2018a (www.mathworks.com) and R version 3.3.2 (http://www.R-project.org). Statistical comparisons were conducted in Matlab including linear mixed models (LinearMixedModel class, fitlme()). Our mixed models included fixed effects of the condition, ROI, and hemisphere as well as a random subject intercept. P-values were obtained by a likelihood ratio test that compares a full model with the effect of interest against a reduced model without this effect (LinearMixedModel class, compare()).

### PS analyses

Pattern similarity (PS) analyses (Kriegeskorte et al., 2008a) were conducted on beta maps generated from unsmoothed data in native subject space as described in (Dimsdale-Zucker et al., 2018) and summarized here. Following the procedure described by Mumford et al (Mumford et al., 2012; Ritchey et al., 2015), single trial models were generated to estimate a unique beta map for every trial in a run (N = 63). Within each single trial model, the first regressor modeled the trial of interest with a stick function, the second regressor modeled all other trials in that run, six regressors were used to capture motion, and any additional spike regressors as identified by our QA scripts were used to capture additional residual variance. Following beta estimation, outlier values were identified by setting a z-scored beta threshold of 0.7–0.85 based on visual inspection of the distribution of z-scored beta values for all subjects. This resulted in an average of 9.87% (mean = 6.22 trials, SD = 7.10 trials) excluded beta values per run for each participant.

Voxel-wise patterns of hemodynamic activity were separately extracted for each ROI from the single trial beta images. Within each ROI, correlations (Pearson’s R) were computed between these trial-wise betas to yield a trial-by-trial correlation matrix that related each voxel’s signal on a trial to all other trials across all runs. Statistical analyses tested for differences in correlations between trial pairs based on encoding location (room and house). Only between-run correlations were used to maximize the number of possible trial pairs without mixing within- and between-run correlations. Trial pairs of interest were extracted from these trial-by-trial correlation matrices. For all conditions, we restricted comparisons to trials in which participants made a correct “remember” response (during MRI scanning), correctly placed the object in the room it was seen in the immediate post-encoding test, and also correctly identified the object’s spatial (house) context (post-MRI spatial source task). To control for the possibility that any observed effects are attributed to temporal context (these effects were previously reported in (Dimsdale-Zucker et al., 2018), we eliminated trial pairs that had been studied within the same video to ensure that any observed effects could uniquely be attributed to spatial context.

Additionally, we used bootstrapping (Westfall and Young, 1993) to validate that the different sizes of groups represented in each bar are not biassing the results. We resampled the larger group 10^3^ times and compared the medians of the two groups. We verified the sign of the difference between medians matches our reported results in more than 99.95% of the bootstrapped groups.

### Data availability

Minimally processed data needed to regenerate analyses and figures can be found online (https://osf.io/5th8r/) as well as relevant analysis code (https://github.com/mgevasagiv/spatcon_2).

## Footnotes

1 We have also conducted statistical analysis using one-way ANOVA which corroborated these results.

2 For ease of visualization, for pattern similarity comparisons we have chosen to plot means with error bars (standard error of the mean) to illustrate the observed data. This does not exactly recapitulate the way the statistical comparisons were computed since these were performed as mixed effects models.

## Notes

https://osf.io/5th8r/

### Competing Interest Statement

The authors have declared no competing interest.

