## Supplemental Figures 1-2 for "Proximity to Boundaries Reveals Spatial Context Representation in Human Hippocampal CA1"

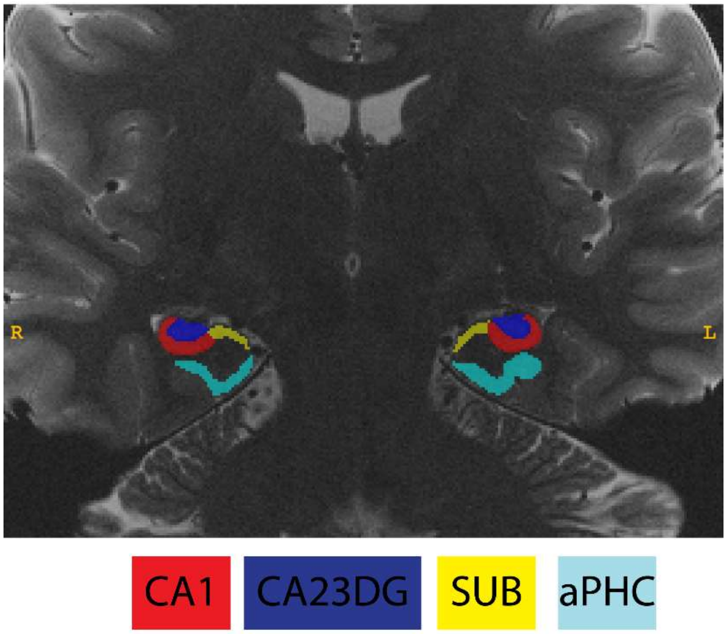


**Supplementary Figure 1:** Segmentations of hippocampal subfields (Cornu Ammonis [CA] 1, a combined CA2, CA3, and dentate gyrus [CA23DG] region, and subiculum [SUB]) and anterior parahippocampal cortex. Segmentations are depicted in the coronal plane of a T2 image (modified from (Dimsdale-Zucker et al., 2022), using the same segmentation protocol).

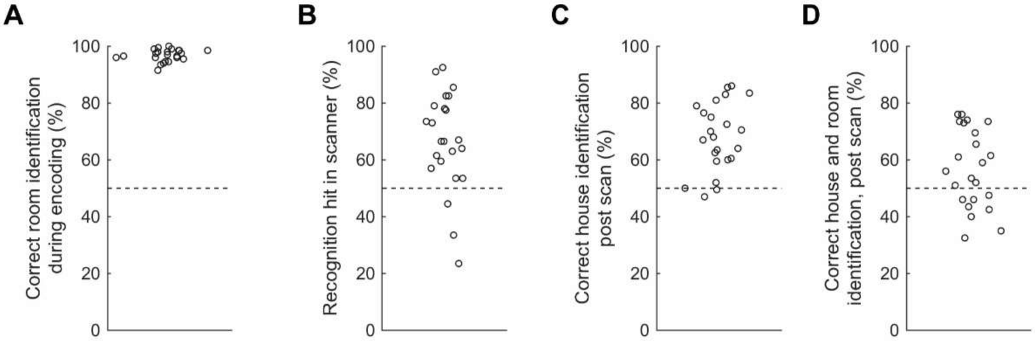


**Supplementary Figure 2: Performance measures across the experimental day**: (**A**) During the encoding phase, subjects were asked to place the objects they have seen in the video on a bird's eye view of the house (Methods, Fig 1B). Performance for this task was extremely high (mean ±std – 96% ±2.1). (**B**) Inside the scanner, subjects were asked to recognize objects they have seen during the encoding phase (Methods), (mean ±std – 66% ±17). (**C**-**D**) Following the scan, subjects were tested in the lab – and asked to recall the house (C) and room (D) for each object learned (Methods), (mean ±std – 68% ±11 and – 56% ±13 for choosing the correct house and room respectively). [Each dot represents the performance of a single subject].
